# Cloning, molecular evolution and functional characterization of ZmbHLHl6, the maize ortholog of OsTIP2 (OsbHLH142)

**DOI:** 10.1101/140806

**Authors:** Yongming Liu, Jia Li, Gui Wei, Yonghao Sun, Yanli Lu, Hai Lan, Chuan Li, Suzhi Zhang, Moju Cao

## Abstract

Abstract Basic helix-loop-helix (bHLH) transcription factors play key roles in plant male reproduction. More than 14 bHLH proteins related to pollen development have been cloned from rice and *Arabidopsis*. However, little is known about the role of the bHLH family in maize microspore development. In this study, the bHLH transcription factor ZmbHLH16 was cloned. ZmbHLH16 shares high similarity with the OsTIP2 (OsbHLH142) protein, a master regulator of the developmental coordination of male reproduction in rice. Expression characterization analysis showed that ZmbHLH16 is preferentially expressed in male reproductive organs and is located in the nucleus. Through nucleotide variation analysis, 36 polymorphic sites in ZmbHLH16, including 23 SNPs and 13 InDels, were detected among 78 maize inbred lines. Neutrality tests and linkage disequilibrium analysis showed that ZmbHLH16 experienced no significant evolutionary pressure. A yeast one-hybrid experiment showed that the first 80 residues in the N-terminus of ZmbHLH16 had transactivation activity, whereas the full length did not. To identify potential ZmbHLH16 interactors, 395 genes that shared similar expression patterns in a genome-wide search were obtained through coexpression analysis. Among these genes, the transcription factor ZmbHLH51 had an expression pattern and subcellular localization similar to those of ZmbHLH16. The interaction between ZmbHLH51 and ZmbHLH16 was verified in yeast cells. In addition to the typical bHLH domain, other regions of ZmbHLH16 were necessary and adequate for its heterodimerization with ZmbHLH51. Our results contribute to a solid foundation for further understanding the functions and mechanisms of ZmbHLH16.

## 1 Introduction

Maize is the most important crop in the world for its utilization in food and industrial materials. At present, there is a rising demand for maize crop yields (Ray et al., 2013). Benefitting from hybrid vigor, male sterility can be used for hybrid maize seed production to increase crop yield. Therefore, the study on male sterility is of great value in application. Till now, several maize genic male sterile (GMS) genes such as *ms8* (Wang et al., 2013), *ms9* (Albertsen et al., 2016), *ms26* (Djukanovic et al., 2013), *ms32* (Moon et al., 2013), *ms44* (Fox et al., 2017) and *ms45* (Albertsen et al., 1993), have been cloned and illuminated for their abortion mechanism. These findings not only contribute to maize heterosis utilization but also expand our understanding of the maize male reproduction. Conventionally, genic male sterile genes are mainly identified through mutant analysis. With the development of gene-editing technology, more male sterile genes are now from the direct editing of some key genes involving pollen development in maize (Svitashev et al., 2015; Mark Cigan et al., 2016; Qi et al., 2016; Svitashev et al., 2016). As a result, the identification of key genes in male reproduction is becoming increasingly important. More than 10,000 genes have been shown to be expressed specifically in maize male fertility development (Ma et al., 2009). Transcription factors (TFs) play key roles in regulating their spatial and temporal-specific expression. Interestingly, TFs might also be the target genes of some small RNAs in plant meiotic processes (Chen, 2004; Alonsoperal et al., 2010; Yu et al., 2013). These above findings indicate that TFs play important roles in plant reproduction. In maize, a total of 2,298 TFs have been identified, and some show tissue-specific expression (Jiang et al., 2012). Key TFs in maize meiosis have been identified using high-throughput techniques such as microarray hybridization and transcriptome sequencing (Dukowic-Schulze et al., 2014a; Dukowic-Schulze et al., 2014b; Zhang et al., 2014). However, only two pollen development-related transcription factors—*ms9* (R2R3-MYB) and *ms32* (basic helix-loop-helix (bHLH))—have been cloned in maize (Moon et al., 2013; Albertsen et al., 2016). To reveal the regulatory mechanism of maize pollen formation, it is imperative to identify additional TFs involved in maize male fertility.

The bHLH proteins compose one of the largest plant transcription factor families. In rice and maize, 178 and 276 bHLH TFs have been identified respectively (Li et al., 2006; Carretero-Paulet et al., 2010; Jiang et al., 2012). Abnormal functions of some bHLH TFs may lead to plant male sterility. In Arabidopsis, ten bHLH proteins related to pollen development have been isolated: AtAMS (Sorensen et al., 2003; Xu et al., 2014), AtDYT1 (Zhang et al., 2006; Feng et al., 2012), AtbHLH10 (Zhu et al., 2015), AtbHLH89 (Zhu et al., 2015), AtbHLH91 (Zhu et al., 2015), AtJAM1 (Nakata et al., 2013), AtJAM2 (Nakata & Ohme-Takagi, 2013), AtJAM3 (Nakata & Ohme-Takagi, 2013), AtMYC5 (Figueroa & Browse, 2015) and AtBIM1 (Xing et al., 2013). Moreover, these male sterile mutants have unique male reproduction-deficient characteristics. For example, the *ams* mutant exhibits total male sterility without any visible pollen; the *dyt1* mutant can produce few pollen grains with a low rate of self-fertility; and the single mutants of *AtbHLH10, AtbHLH89*, and *AtbHLH91* develop normally, with only their various double and triple combinations defective in pollen development (Sorensen et al., 2003; Li et al., 2006; Zhu et al., 2015). These differences in male sterile characteristics might result from the functional divergence of bHLH TFs. Tapetal cells provide energy and nutrition for microspore development via programmed cell death (PCD) at appropriate anther developmental stages (Zhang et al., 2008). In rice, bHLH TFs including OsUDT1 (Jung et al., 2005), OsTDR (Li et al., 2006; Zhang et al., 2008), OsEAT1/OsDTD1 (Ji et al., 2013; Niu et al., 2013), and OsTIP2 (OsbHLH142) (Fu et al., 2014; Ko et al., 2014) have been found to be essential for anther tapetal cell development. These above rice bHLH TF dysfunctions lead tapetal cells to undergo abnormal PCD, thereby causing complete male sterility. In conclusion, all above show that bHLH TFs play important roles in regulating stamen development. At present, the study about bHLH TFs related to maize male fertility is few, there is a need to characterize more bHLH proteins in maize male reproduction.

*TIP2* (OsbHLH142) acts as a key regulator of tapetum development in rice (Fu et al., 2014; Ko et al., 2014). The *tip2* mutant displays complete male sterility, with three undifferentiated anther wall layers (epidermal, fibrous and middle layer) and abortion of tapetal programmed cells death(Fu et al., 2014). In this study, the transcription factor ZmbHLH16, which is homologous to OsTIP2 (OsbHLH142), was isolated from maize. Its structure, phylogeny, expression and subcellular localization, molecular evolution, and regulatory characteristics were investigated.

## 2 Materials and methods

### 2.1 Plant materials

Spikelets from maize inbred line A619 were collected for ZmbHLH16 (GRMZM2G021276_T02) and ZmbHLH51 (GRMZM2G139372_T07) CDS (coding sequence) cloning. Ears, main stalks, stems, and spikelets were taken from maize inbred A619 for ZmbHLH16 expression analysis. Seeds from 78 inbred lines (see Supplementary Table S1) were used to amplify the genome sequence of ZmbHLH16.

### 2.2 DNA and RNA extraction

Genomic DNA was extracted from seeds using a modified cetyltrimethylammonium bromide (CTAB) method (Porebski et al., 1997). Total RNAs were isolated from the above frozen samples with TRIzol reagent (Takara, China) and DNase I to eliminate any genomic DNA. One microgram of total RNA from each sample was used to synthesize cDNA via the PrimeScript™ RT Reagent Kit (Takara, China).

### 2.3 CDS cloning of ZmbHLH16 and phylogenetic analysis

BlastP^1^ was used to identify OsTIP2 (OsbHLH142) homologous genes in the maize genome (B73 assembly v3) with the amino acid sequence of OsTIP2 (OS01G0293100). The CDS of ZmbHLH16 was amplified from cDNA templates of A619 spikelets with the following primers: 5′-ATGTATCACCCGCAGTGCGAGCT-3′ and 5′-TGTACTCGTCCACCACTTCCAT-3′. High-fidelity KOD FX (Toyobo, Japan) was used for gene cloning according to the manufacturer′s instructions. The purified PCR products were inserted into the pEASY Blunt Simple cloning vector (TransGen, China) and sequenced by Tsingke Biotech with an ABI 3730XL DNA Analyzer. The ZmbHLH16 amino acid sequence was acquired based on amplifying its CDS from A619 using the online program SoftBerry FGENESH^2^, and its conserved domain was predicted using the NCBI CD tool^3^. Then, 16 bHLH TFs that were reported to be involved in microspore development were retrieved from Gramene^4^ to construct a phylogenetic tree using the neighbor-joining method with MEGA v5.10 (Kumar et al., 2008), and the robustness of the findings was verified via 1000 bootstrap replicates. The accession numbers of the 16 TFs are as follows: AtAMS (AT2G16910), AtDYT1 (AT4G21330), AtbHLH10 (AT2G31220), AtbHLH89 (AT1G06170), AtbHLH91 (AT2G31210), AtJAM1 (AT2G46510), AtJAM2 (AT1G01260), AtJAM3 (AT4G16430), AtMYC5 (AT5G46830), AtBIM1 (AT5G08130), OsUDT1 (OS07G0549600), OsTDR1 (OS02G0120500), OsEAT1 (OS04G0599300), OsTIP2 (OS01G0293100), ZmMS32 (GRMZM2G163233) and SlMS1035 (Solyc02g079810).

### 2.4 Molecular evolution analysis of ZmbHLH16

The genomic sequences of ZmbHLH16, including its 5′ and 3′ untranslated regions (UTRs), were amplified from 78 maize inbred lines (see Supplementary Table S1 for details) with the primers 5′-GGAAGGAGGAAACCAAGTCG-3′ and 5’-TGTAACGAGCAAGCGGATTTA-3′. PCR was performed according to the manufacturer’s protocol using high-fidelity polymerase KOD FX (Toyobo, Japan). PCR-amplifying fragments were purified and sequenced directly using an ABI 3730XL DNA Analyzer manufactured by Tsingke Biotech. After ambiguous sequences were manually deleted, the sequence polymorphisms of ZmbHLH16 among the 78 maize inbred lines were analyzed using CodonCode Aligner 6.0.2 software (CodonCode Corporation, Dedham, MA). For molecular evolution analysis, certain parameters were calculated as follows: (1) the nucleotide diversity of common pairwise nucleotide difference per site (π) with DnaSP 5.0 (Librado & Rozas, 2009); (2) in neutrality tests, the evolutionary pressure in ZmbHLH16 via Tajima’s D test (Tajima, 1989) and Fu and Li’s statistics (Fu & Li, 1993); (3) the LD matrix of ZmbHLH16 was characterized by evaluating *r*^2^ values based on SNPs and InDels (minor allele frequency (MAF) ≥0.05) in TASSEL 2.0 (Bradbury et al., 2007). An LD plot was obtained in Haploview 4.2 (Barrett et al., 2005), and the LD decay was assessed by averaging *r*^2^ values with a distance of 250 bp.

### 2.5 Transactivation activity analysis of truncated ZmbHLH16

The ZmbHLH16 CDS contains 1098 bp encoding a protein with 365 amino acids. To investigate its transcriptional activating ability and retain its conserved bHLH domain, the ZmbHLH16 CDS sequence was divided into four parts; the first three parts each contained 240 bp (labeled A ^1–80 a.a.^, B ^81–160 a.a.^· and C ^161–240 a.a.^), and the last part contained 375 bp (labeled D ^241–365 a.a.^) (Fig 3A). Three new units—ZmbHLH16 (E) ^1–160 a.a.^, (F) ^81–240 a.a.^, and (G) ^161–365 a.a.^—were constructed using combinations of two neighboring parts. The above seven parts, termed ZmbHLH16 (A)-(G), were artificially synthesized, and sequencing confirmation was performed by Tsingke Biotech. In total, eight fragments, including ZmbHLH16 CDS and ZmbHLH16 (A)-(G), were individually inserted into the pGBKT7 vector using the In-Fusion cloning method (Vazyme ClonExpress II One Step Cloning Kit, Vazyme Biotech Co., Ltd, China) to analyze their transcriptional activation activity (see Supplementary Table S2 for all primers used in the experiment). All recombinant pGBKT7 vectors were transformed into AH109 yeast strains (TIANDZ, China) via the lithium acetate-mediated approach. The transformants were cultivated on SD/-Trp medium for 2–3 days at 28°C. Bacterial PCR was used to identify positive clones. The positive clones were further cultured on SD/-His-Trp medium containing 50 mg/L χ-α-gal (Coolaber, China) for 2–4 days at 28°C to test their transactivation activity. The pGBKT7 and pGBKT7-GAL4 AD vectors were used as negative and positive controls, respectively.

### 2.6 Coexpression analysis and identification of maize male reproduction-related genes

For coexpression analysis, expression data of genome-wide maize genes in 20 tissues and 66 periods were obtained from q-teller^5^, and the Pearson correlation coefficients (PCCs) between ZmbHLH16 and other genes were calculated. Cluster3.0 (de Hoon et al., 2004) was used for target gene (PCC>0.6) cluster analysis based on Euclidean distance and complete linkage. A heatmap was drawn using Java Treeview (Saldanha, 2004). Next, to gain deeper insight into the molecular mechanism underlying ZmbHLH16, all target genes (PCC>0.6) were queried with E-value<1e^−5^ in the Plant Male Reproduction Database^6^, which contains 548 male-fertility-related genes in Arabidopsis. All maize gene sequences were retrieved from MaizeGDB^7^. To characterize the putative function of ZmbHLH16-coexpressed genes, GO terms for all target genes (PCC>0.6) were taken from AGRIgo^8^, and GO enrichment analysis was performed using OmicShare tools^9^.

### 2.7 Expression characteristics of ZmbHLH16 and ZmbHLH51

The primers for the semi-quantitative expression analysis of ZmbHLH16 were 5′-CCTCATGCACCTCATACC-3′ and 5′-CAGCTCCTGGATGTACTC-3′. At the same time, the primer sequences 5′-CTGGAGGTCACCAACGTCAA-3′ and 5′-AGCGAGTCCCTCAGTCTGTC-3′ were used for ZmbHLH51 expression analysis. The 18S gene in this experiment was used as the internal control, and its amplifying primers were 5′-CTGAGAAACGGCTACCACA-3′ and 5′-CCCAAGGTCCAACTACGAG-3′ (Hu et al., 2011).

The localization patterns of the ZmbHLH16 and ZmbHLH51 proteins were investigated using transient transformation in rice protoplasts. For ZmbHLH51 CDS cloning, we first amplified the cDNA sequence with the primers 5′-GAGCAGTGATGTGAATTGCG-3′ and 5′-TCAAGCGAGGTATTGGAGGA-3′ using high-fidelity KOD FX polymerase from A619 and inserted it into the pEASY blunt-cloning vector. The CDSs of ZmbHLH16 and ZmbHLH51 lacking the stop codons were individually fused to the N-terminus of enhanced green fluorescent protein (eGFP) in pCAMBIA2300-P_35S_ by subcloning using the In-Fusion cloning method. Two recombinants, pCAMBIA2300-P_*35S*_:ZmbHLH16-eGFP and pCAMBIA2300-P_*35S*_:ZmbHLH51-eGFP, were constructed to assess the localization of these proteins. The empty pCAMBIA2300-P35S-eGFP vector was used as a control in this experiment. The recombinant vectors pCAMBIA2300-P_*35S*_:ZmbHLH16-eGFP and pCAMBIA2300-P_*35S*_:ZmbHLH51-eGFP and the control vector were transformed into rice protoplasts using polyethylene glycol (PEG), as described previously (Bart et al., 2006). The green signals (Ex=488 nm, Em=507 nm) were detected using a TCS-SP8 fluorescence microscope (Leica, Germany).

### 2.8 Protein–protein interactions

To confirm the interaction between ZmbHLH16 and ZmbHLH51, a yeast two-hybrid assay was conducted. The CDSs of ZmbHLH16 and ZmbHLH51 were inserted into the pGBKT7 and pGADT7 vectors, respectively. The pGBKT7-ZmbHLH16 vector without autoactivation activity was constructed as above. The pGADT7-ZmbHLH51 vector was constructed using the In-Fusion cloning method by subcloning from pCAMBIA2300-P_*35S*_:ZmbHLH51. The recombinant vectors pGBKT7-ZmbHLH16 and pGADT7-ZmbHLH51 were co-transformed into AH109 yeast competent cells according to operating instructions. The transformants were cultivated on SD/-Leu-Trp medium at 28°C for 2–3 days, and positive clones were confirmed using PCR. Positive clones were further cultured on SD/-Ade-His-Leu-Trp medium containing 50 mg/L χ-α-gal at 28°C for 2–3 days. The vectors pGBKT7-T and pGBKT7-Lam were used as positive and negative controls, respectively. After confirming the interaction between ZmbHLH16 and ZmbHLH51, to investigate the interaction domain in ZmbHLH16, regions of ZmbHLH16 without autoactivation activity were inserted into the bait vector pGBKT7 and then co-transformed with the prey vector pGADT7-ZmbHLH51 into the AH109 yeast competent cells.

## 3 Results

### 3.1 ZmbHLH16 is a typical bHLH transcription factor

TIP2 is a key regulator involved in rice anther development. BlastP analysis showed that the gene model GRMZM2G021276 (ZmbHLH16) had the highest homology score with OsTIP2 (OsbHLH142). The ZmbHLH16 CDS was obtained from the maize inbred line A619. Further analysis showed that the ZmbHLH16 coding sequence contained 1098 bp encoding a protein of 365 amino acids that had 66.85% amino acid sequence identity with OsTIP2 (OsbHLH142). Analysis with NCBI CD software revealed that the amino acid sequence of ZmbHLH16 included a typical bHLH DNA-binding domain (Fig 1A). The bHLH interaction and function (BIF) domain participates in bHLH protein localization, transcriptional activity and dimerization (Cui et al., 2016). The BIF domain was also found in the C-terminal (288–363 a.a.) region of ZmbHLH16. Furthermore, phylogenetic analysis of 17 bHLH transcription factors related to microspore development illustrated that these bHLH TFs were highly conserved, with most bootstrap values >90%, and that ZmbHLH16 was the most closely related to OsTIP2 (Fig 1B). Together, the above results indicated that ZmbHLH16 protein included the typically conserved domain of bHLH TFs and might play a crucial role in male reproduction in maize.

**Fig 1.**
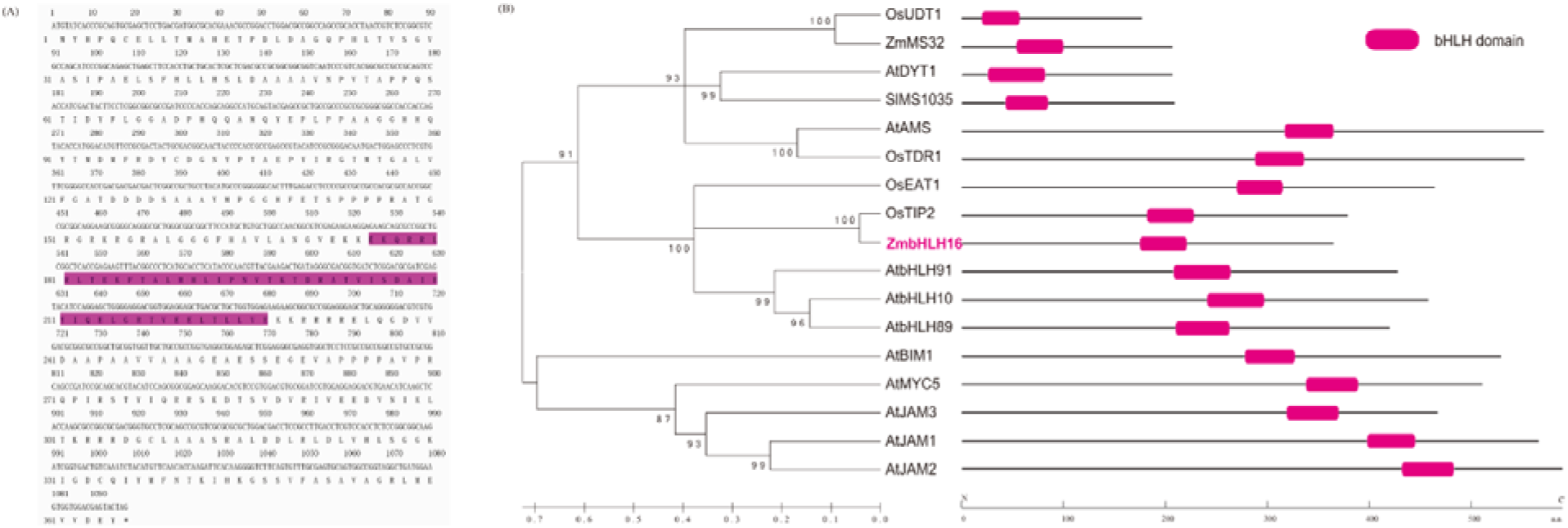
Structure and phylogenetic analysis of ZmbHLH16. (**A**) Nucleotide and deduced amino acid sequences of ZmbHLH16 CDS. Shaded regions are the conserved DNA-binding domains of the bHLH protein. Bold letters show conserved tryptophan residues in the bHLH domain. (**B**) Phylogenetic relationship of ZmbHLH16 and other bHLH proteins related to microspore development. The two scale bars indicate branch length and amino acid length.

### 3.2 ZmbHLH16 is highly evolutionarily conserved in maize germplasm

To analyze its molecular evolution, the DNA sequences of ZmbHLH16 were amplified from 78 maize inbred lines. The ZmbHLH16 genomic sequence is divided into seven regions with a length of 2,514 bp (Table 1). Based on the MAF≥0.05, 36 polymorphism sites within ZmbHLH16 were identified, including 23 SNPs and 13 InDels, with one SNP/InDel per 109/193 bp (Table 1). Among 23 SNPs, 13 (57%) and 10 (43%) involved transitions and transversions, respectively. Further analysis showed that there were 4 amino acid variations in ZmbHLH16 among 78 inbred lines, with 3 mutations in the first exon and the fourth mutation in the second exon. Comparison analysis showed that nucleotide variations were not evenly distributed in ZmbHLH16, and introns had higher sequence diversity (3.1 polymorphisms/100 bp) than UTRs (1.24 polymorphisms/100 bp) and exons (1.18 polymorphisms/100 bp). Moreover, a nucleotide polymorphism test in DnaSP showed the highest nucleotide diversity ratio (π=6.15×10^−3^) in the first intron but no significant nucleotide variation in the 3rd intron or 3′-UTR (Table 2).

**Table 1.**
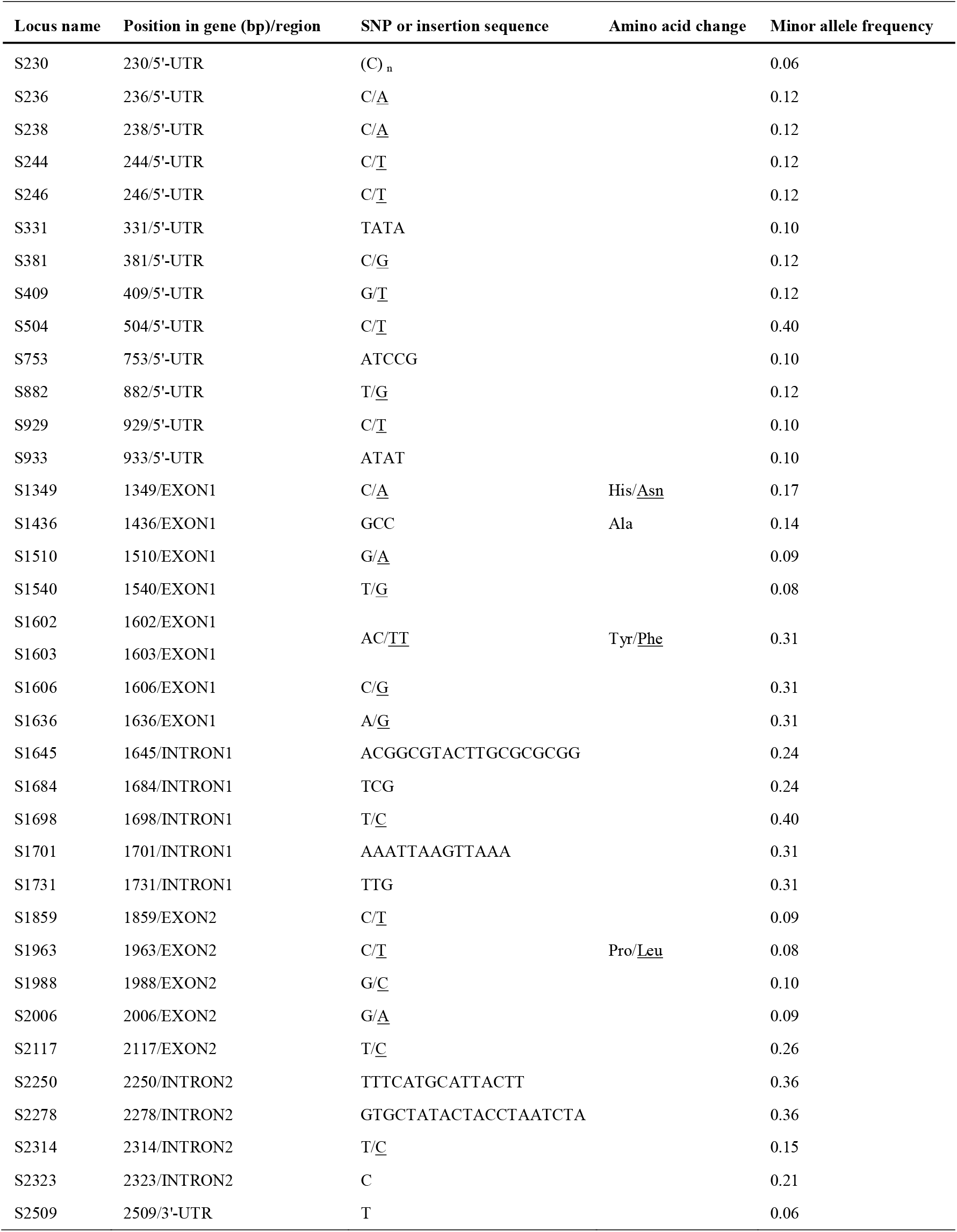
Single-nucleotide polymorphisms (SNPs) and insertion-deletion polymorphisms (InDels) of ZmbHLH16 among 78 maize inbred lines. Underlined letters represent minor alleles.

To investigate whether ZmbHLH16 has experienced selection pressure, various regions of ZmbHLH16 were assessed (Table 2). In Tajima’s D test and Fu and Li’s test, no significant difference was obtained across all regions of ZmbHLH16. This result indicated that ZmbHLH16 experienced no significant selective pressure and underwent neutral selection. Elevated LD is usually expected for genes under selection (Bomblies & Doebley, 2005). Thus, to further confirm whether ZmbHLH16 underwent directional selection, its LD patterns and LD decay were also calculated. In the LD matrix, no obvious LD block was found in the ZmbHLH16 genome sequence (Fig 2A). A schematic diagram of LD decay represented by plots of *r*^2^ showed that the LD level dropped to 0.1 at approximately 1,300 bp (Fig 2B), indicating a more rapid decay rate than the average 1.5 kb of several genes under selection in maize (Remington et al., 2001). Therefore, our above results also reflected the conserved evolution of ZmbHLH16 in the maize germplasm.

**Fig 2.**
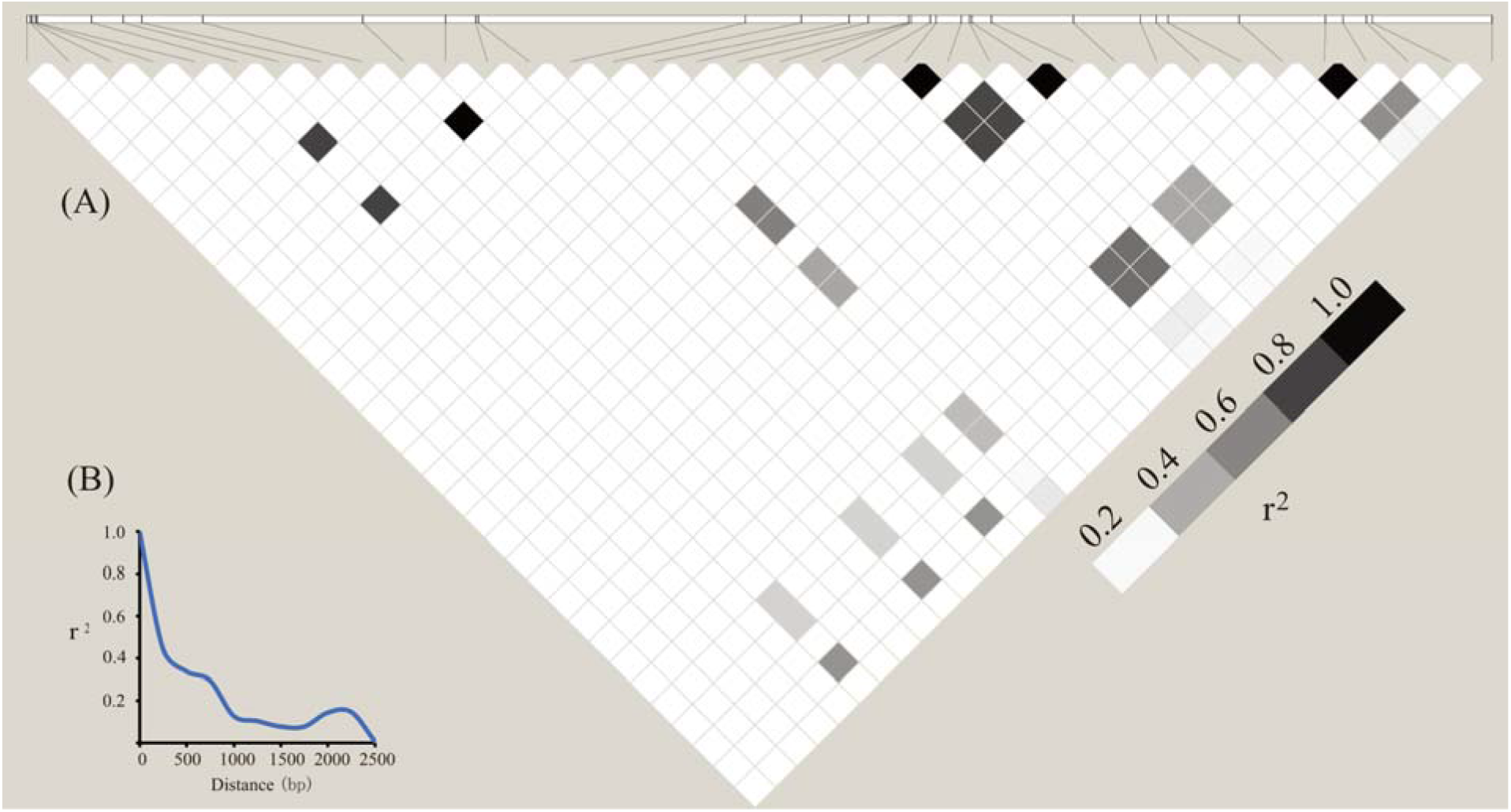
LD block and LD decay pattern of ZmbHLH16. (**A**) Matrix of pairwise LD of DNA polymorphisms (MAF≥0.05) in ZmbHLH16. The shaded boxes indicate the LD standard (*r*^2^). (B) LD decay in the DNA sequence of ZmbHLH16 in maize. The *x* axis represents the distance between polymorphic sites, and the *y* axis represents the average *r^2^* value for each distance category (250 bp).

**Table 2.** ZmbHLH16 nucleotide diversity and neutrality test. Due to a lack of significant polymorphism sites (MAF≥0.05), values for neutrality tests in Exon 3 and the 3′-UTR are not given.

| Region | 5'-UTR | Exon 1 | Intron 1 | Exon 2 | Intron 2 | Exon 3 | 3'-UTR | Overall |
| --- | --- | --- | --- | --- | --- | --- | --- | --- |
| $\pi(\times 10^{-3})$ | 1.85 | 4.13 | 6.15 | 2.48 | 2.39 | 0 | 0 | 2.58 |
| Tajima's D | -0.65 | 0.48 | 0.42 | -0.28 | -0.91 | / | / | -0.26 |
| Fu andLi's D | -0.92 | -0.64 | -1.01 | 0.22 | -1.90 | / | / | -1.21 |
| Fu andLi's F | -0.98 | -0.30 | -0.67 | 0.06 | -1.86 | / | / | -1.01 |
| Length (bp) | 1042 | 597 | 121 | 432 | 168 | 72 | 82 | 2514 |

### 3.3 Only the N-terminal first 80 residues of ZmbHLH16 have transactivation activity

To identify the activating function of ZmbHLH16, eight fragments of ZmbHLH16 were analyzed in yeast (Fig 3A). Yeast cells with pGBKT7-ZmbHLH16(A) ^1–80 a.a.^ or pGBKT7-ZmbHLH 16(E) ^1–160a.a.^ grew normally on both SD/-Trp and SD/-His-Trp selective media and turned the indicator blue. However, the other six yeast transformants with ZmbHLH16 could only live on the SD/-Trp medium (Fig 3B). In comparison with the living conditions of the transformants containing ZmbHLH16 (A) ^1–8a.a.^, (B) ^81–16a.a.^ and (E) ^1–160a.a.^, this finding suggested that the N-terminal first eighty amino acids of ZmbHLH16 possessed transcriptional activation activity. Based on a comparison with the whole coding region (1–365 a.a.) and region (E) ^1–160 a.a.^, it was inferred that domain (G) ^161–365 a.a.^ of ZmbHLH16 might inhibit its transcriptional activation activity. However, the deficiency of transactivation activity of the full-length ZmbHLH16 might also be due to its poor expression in yeast. In conclusion, the above results indicated that the first 80 amino acids in the N-terminus of bHLH16 could activate transcription in yeast, whereas the full-length version did not.

**Fig 3.**
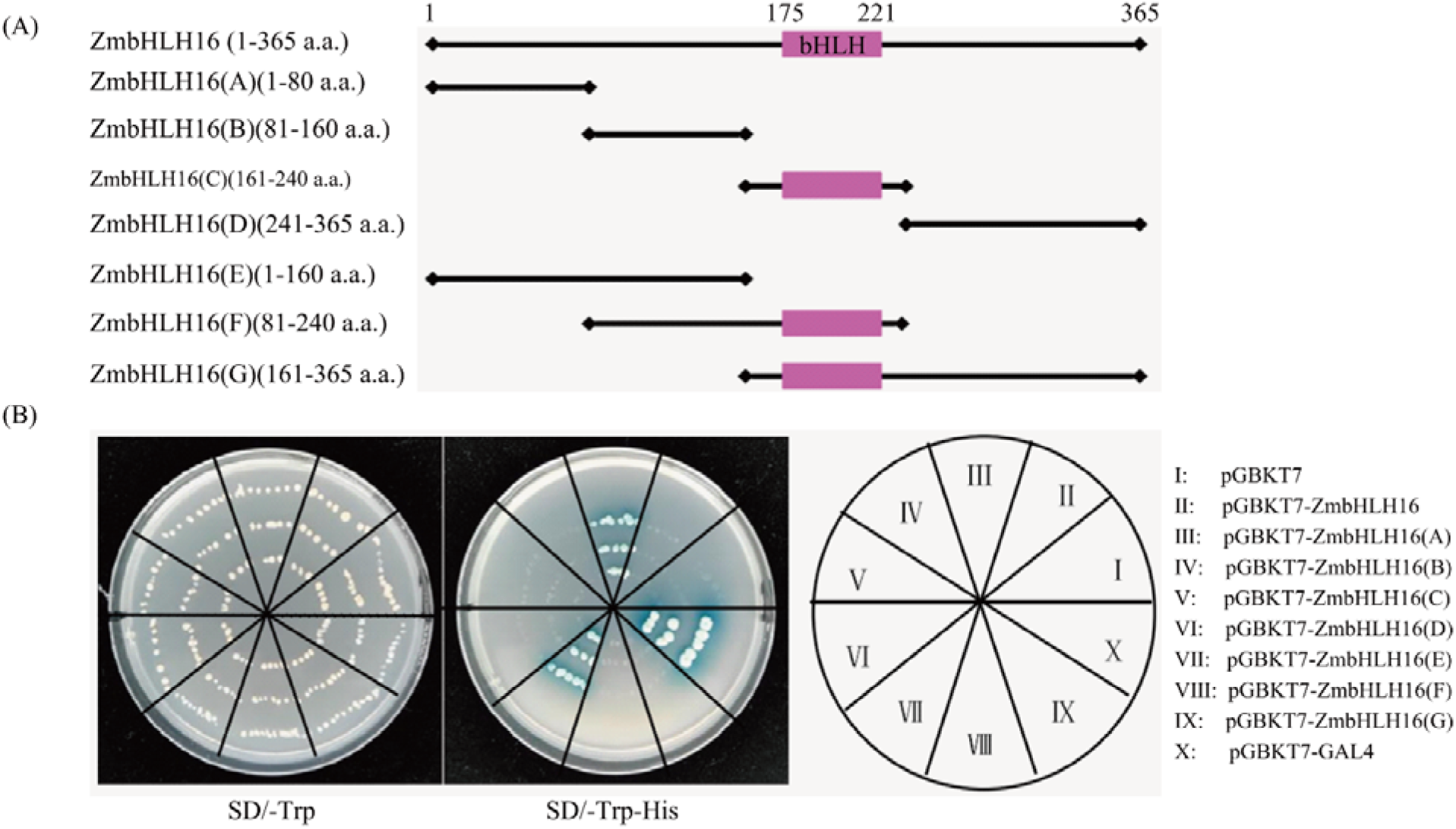
Transactivation activity assays of ZmbHLH16 in yeast. (**A**) Diagram of ZmbHLH16 activation domain. (**B**) Growth of yeast containing various fragments of ZmbHLH16 on selective media (SD/-Trp and SD/-Trp-His).

### 3.4 ZmbHLH16 coexpresses with many male reproduction-related genes

Functionally associated genes are more likely to share similar expression patterns (Fu & Xue, 2010). Coexpression analysis was therefore conducted to identify potential ZmbHLH16 cooperators. A total of 395 maize genes with calculated PCC values above 0.6 were found (Fig 4A and Supplementary Table S3). Among them were three male sterile genes, including *ms8* (GRMZM2G119265), *ms26* (GRMZM2G091822) and *ms44* (AC225127.3_FG003), which shared expression PCC values of 0.9937, 0.8375 and 0.9964 with ZmbHLH16, respectively. In a search of the PMRD database, in maize whole genome we identified 2405 (6.0%) genes showing homologous to Arabidopsis male fertility genes (Supplementary Table S4). In comparison, a higher ratio of 8.8% (35/395) at a significant level *χ*^2^=4.92, *p*=0.0265) was got when ZmbHLH16 coexpressed genes were queried (Table 3). The similar expression pattern to a number of plant reproduction-related genes indicated that ZmbHLH16 might be closely associated with maize male fertility.

**Fig 4.**
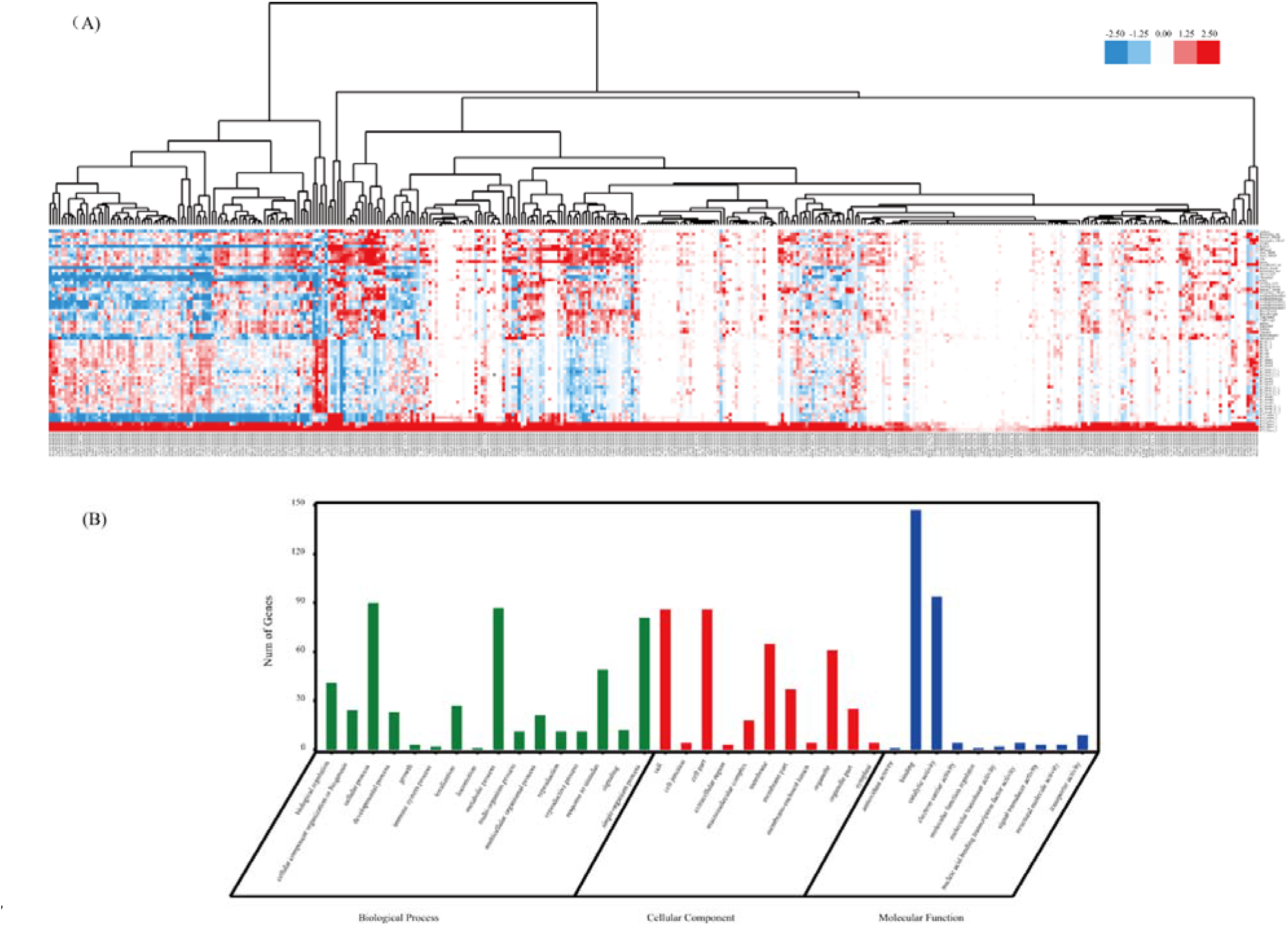
Expression patterns and GO annotations of ZmbHLH16 coexpressed genes. (**A**) A cluster of 395 coexpressed genes based on expression characteristics. Gene expression data were downloaded from q-teller (http://www.qteller.com/qteller3/index.php); the bar indicates the relative gene expression level, which was log_2_-normalized (original data+1). (**B**) GO analysis of 395 coexpressed genes.

**Table 3.**
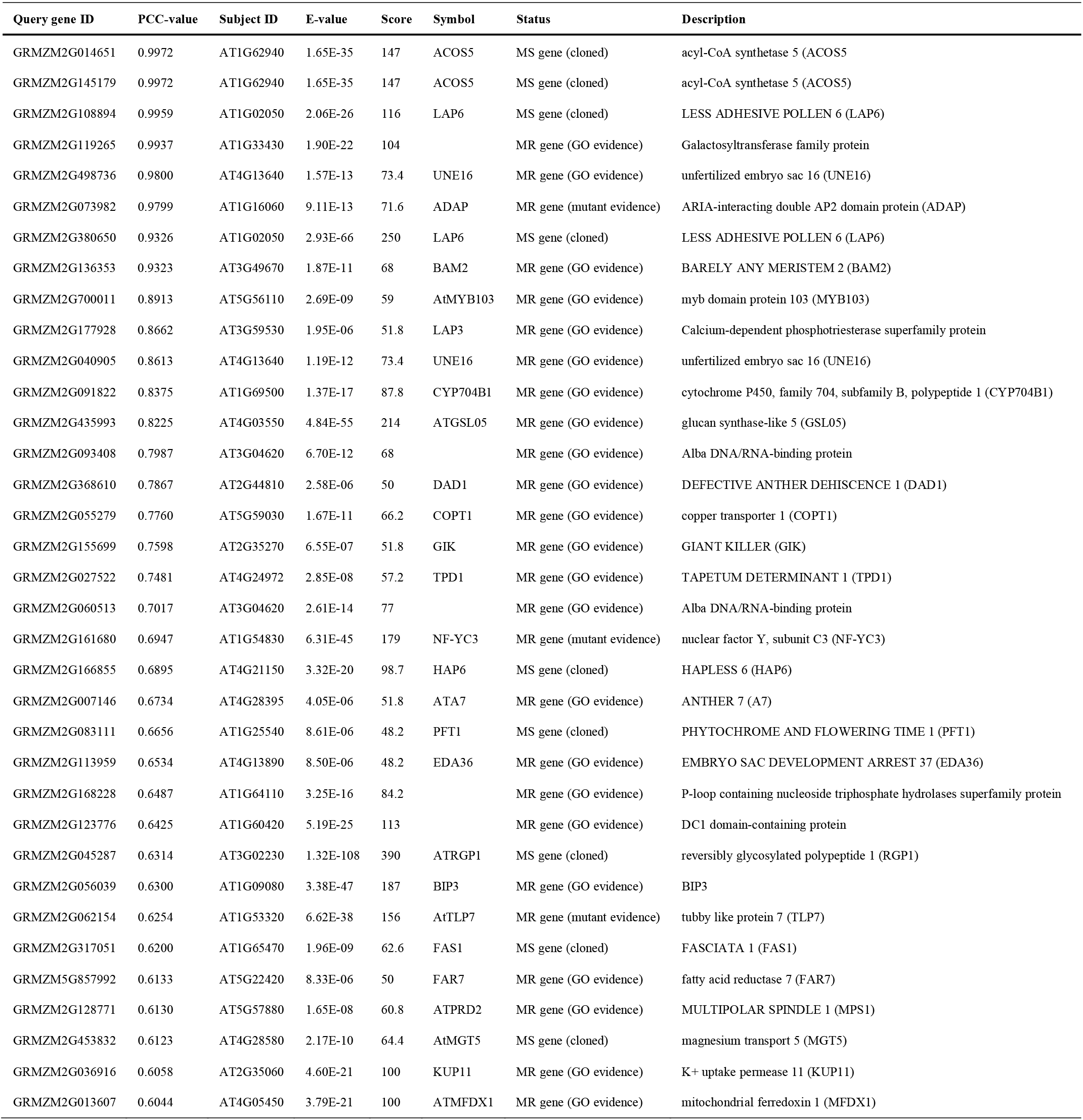
ZmbHLH16 coexpressed genes homologous to Arabidopsis male-sterility (MS)/reproduction (MR) genes.

Next, the 395 coexpressed genes were subjected to GO term analysis (Fig 4B and Supplementary Table S3). In the cellular component, 155 GO terms were enriched and most of these genes were categorized under cells, cell parts, membranes and organelles. For the molecular function category, binding and catalytic activity were the most abundant subcategories. Similarly, previous reports have confirmed that the binding activity and catalytic activity functions are essential for alterations in male fertility (Qu et al., 2015; Zhu et al., 2015; Mei et al., 2016). For biological processes, there were 780 enriched GO terms and cellular processes, metabolic processes, and single-organism processes were the most abundant clusters. Through hypergeometric test at the 0.05 significance level, it was found that some enriched Go terms such as pollen wall assembly, pollen exine formation, pollen development, gametophyte development reached to significant levels compared with the maize background (Supplementary Table S3). These above results supported that ZmbHLH16 might participate in maize pollen formation. Moreover, in the reproduction GO term (GO:0000003), a bHLH transcription factor family member, ZmbHLH51 (GRMZM2G139372), was found, which shared a PCC score of 0.8990 with ZmbHLH16. ZmbHLH51 was homologous to the male sterile gene OsTDR. Accordingly, ZmbHLH51 might be an important factor in maize pollen development. Some studies have indicated that the interactions among bHLH TFs are important for pollen development (Niu et al., 2013; Zhu et al., 2015). Therefore, we next aimed to analyze the interaction between ZmbHLH16 and ZmbHLH51.

### 3.5 ZmbHLH16 and ZmbHLH51 have similar expression characteristics

The expression patterns of ZmbHLH16 and ZmbHLH51 were simultaneously analyzed using semi-quantitative PCR for reproductive and vegetative organs. Both ZmbHLH16 and ZmbHLH51 showed a higher expression level in spikelets than organs (Fig 5A). This finding indicated that ZmbHLH16 and ZmbHLH51 might be closely associated with maize male fertility.

**Fig 5.**
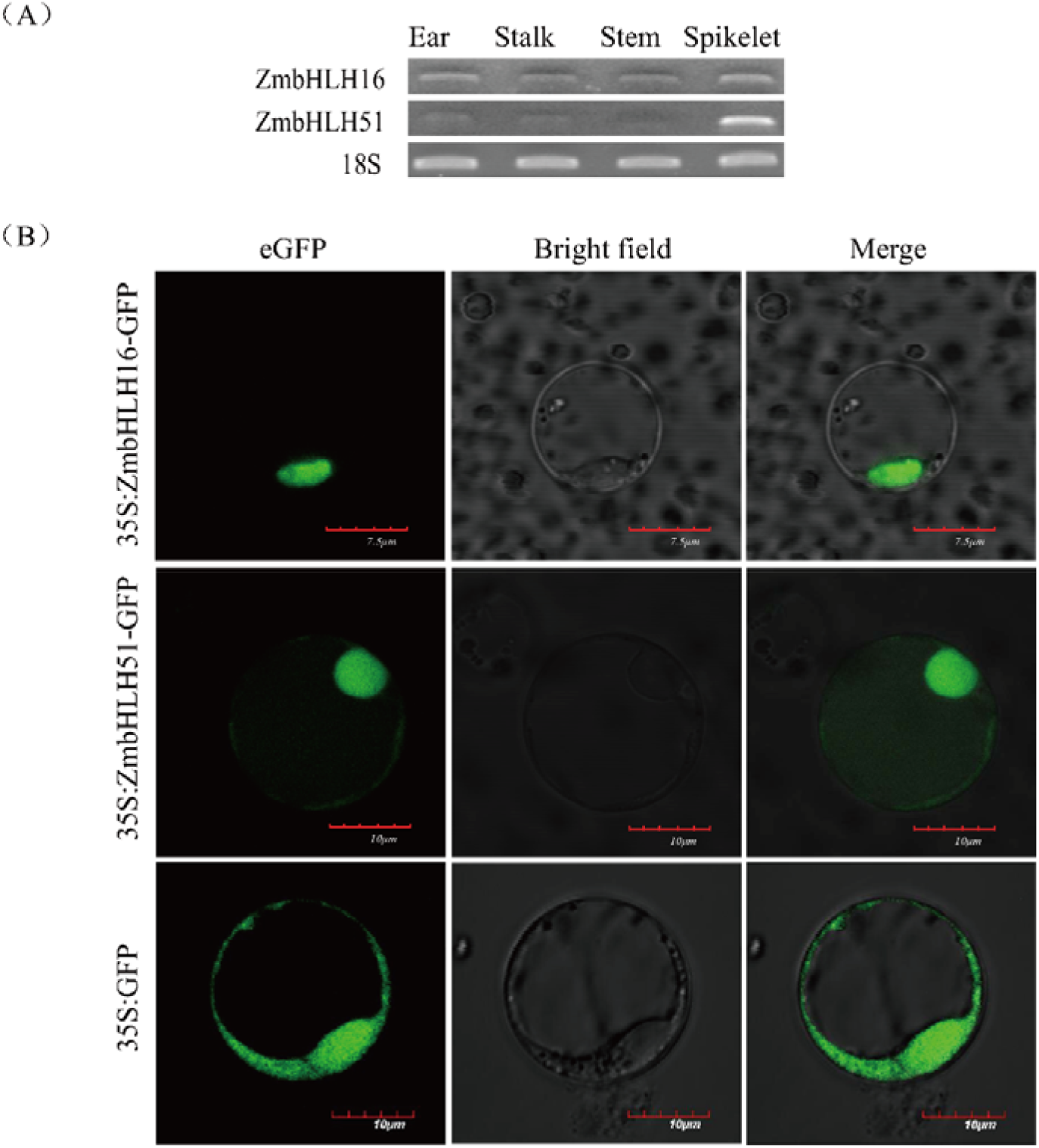
Expression characteristics of ZmbHLH16 and ZmbHLH51. (**A**) Semi-quantitative analysis of ZmbHLH16 and ZmbHLH51 in various organs. The expression of 18S was taken as the reference. (**B**) Subcellular localization analysis of ZmbHLH16 and ZmbHLH51 in rice protoplasts.

Based on the above results, ZmbHLH51 is homologous to the male sterile gene *OsTDR* and might interact with ZmbHLH16. Therefore, the subcellular localization of both ZmbHLH16 and ZmbHLH51 was investigated. As depicted in Fig 5B, the recombinant fusion proteins ZmbHLH16-eGFP and ZmbHLH51-eGFP were only located in the nucleus, whereas the control eGFP was localized to both the cytoplasm and the nucleus. The similar expression profiles and protein localization patterns between ZmbHLH16 and ZmbHLH51 supported their interaction.

### 3.6 ZmbHLH51 interacts with ZmbHLH16

Because the aforementioned results indicated that the two bHLH TFs ZmbHLH16 and ZmbHLH51 had similar expression characteristics and subcellular localization patterns, a yeast two-hybrid assay was used to verify the interaction between the two proteins. As shown in Fig 6A, yeast cells containing pGBKT7-ZmbHLH16 and pGADT7-ZmbHLH51 were able to grow on both SD/-Leu-Trp and SD/-Ade-His-Leu-Trp media, similar to the positive control. The above results confirmed the interaction between ZmbHLH16 and ZmbHLH51.

**Fig 6.**
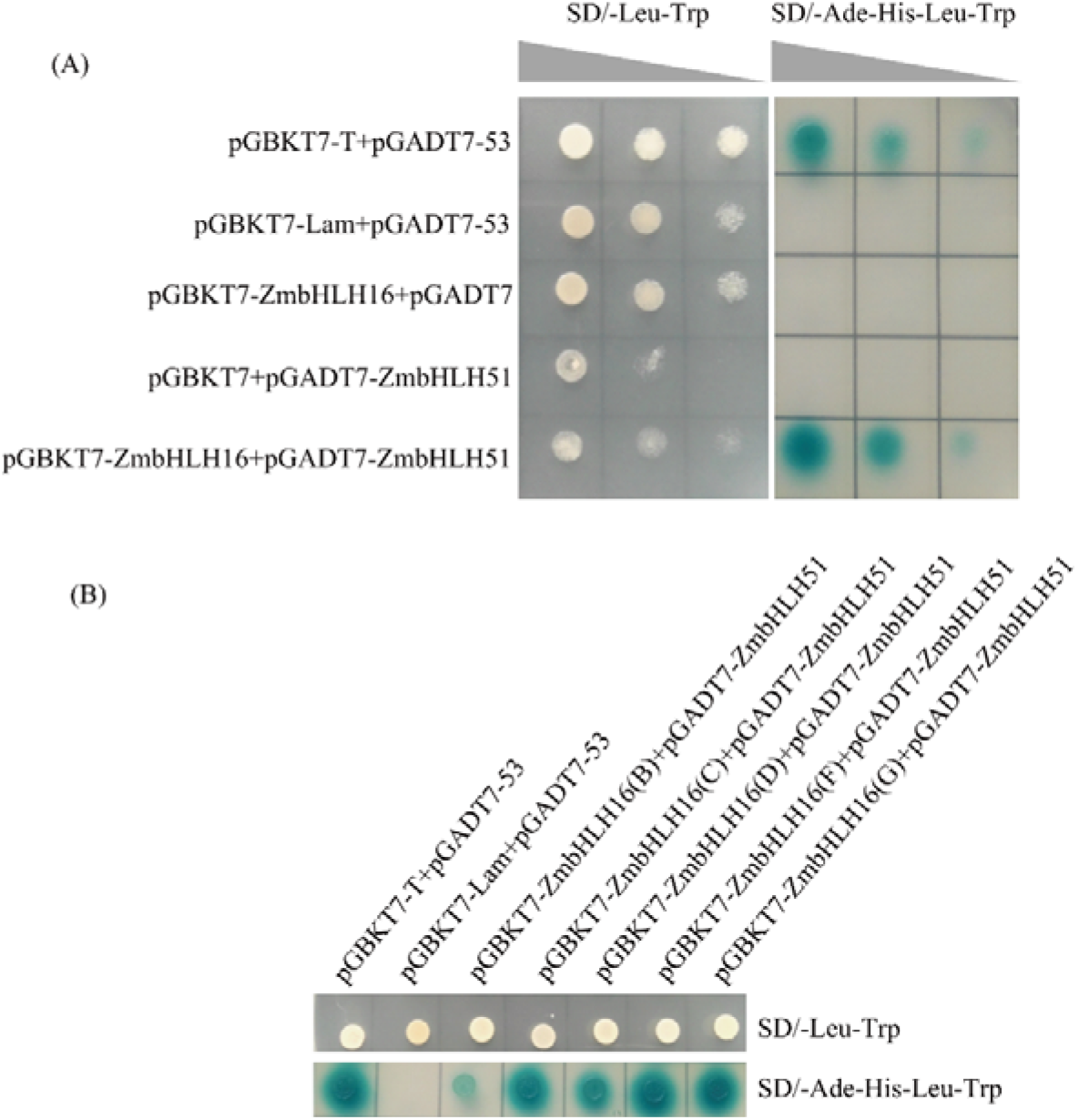
Determination of the interaction between ZmbHLH16 and ZmbHLH51 using yeast two-hybrid analysis. (**A**) Yeast two-hybrid analysis of the interaction between ZmbHLH16 and ZmbHLH51 proteins. Each transformant was stained in media with three relative concentrations (1, 0.1, 0.01) from left to right. (**B**) Yeast two-hybrid mapping of domains involved in the ZmbHLH16-ZmbHLH51 interaction. Regions without transcriptional activation activity, including ZmbHLH16(B)^81–160 a.a.^. (C) ^161–240 a.a.^, (D) ^241–365 a.a.^, (F) ^81–240 a.a.^ and (G) ^161–365 a.a.^, were chosen for analysis.

To map the domains involved in the ZmbHLH16-ZmbHLH51 interaction, fragments without transcriptional activation activity, including ZmbHLH16 (B) ^81–160 a.a.^, (C) ^161–240 a.a.^, (D) ^241–365 a.a.^, (F) ^81–240 a.a.^ and (G) ^161–365 a.a.^, were further analyzed using yeast two-hybrid assays. The conserved bHLH domain is reported to participate in protein homo- or heterodimerization (Pires & Dolan, 2010). As expected, the regions containing the bHLH domain, i.e., (C) ^161–240 a.a.^, (F) ^81–240 a.a.^ and (G) ^161–365 a.a.^, could grow normally on SD/-Ade-His-Leu-Trp media and turned the media blue (Fig 6B). Interestingly, transformants (B) and (D), which lacked the bHLH domain, also survived on the SD/-Ade-His-Leu-Trp synthetic dropout medium. These results indicated that not only the bHLH domain but also other regions in ZmbHLH16 were sufficient and necessary for its heterodimerization with ZmbHLH51.

## 4 Discussion

Male reproduction is a complicated process in plants that involves thousands of genes and many biological processes (Dukowic-Schulze & Chen, 2014; Zhou & Pawlowski, 2014; Rutley & Twell, 2015). Some genes involved in plant anther and pollen development are conserved in plants (Gómez et al., 2015). This phenomenon provides the possibility of elucidating key genes in other species based on homology analysis. Thus, here, we isolated ZmbHLH16 based on homology cloning from OsTIP2, which has been reported to be a master regulator of pollen formation.

In this study, the molecular evolution of ZmbHLH16 was investigated. In the analysis of selective pressure, no significant signal was found in ZmbHLH16 according to Tajima’s D and Fu and Li’s tests. Moreover, a lower nucleotide diversity ratio (π=2.58×10^−3^) was observed in all regions of ZmbHLH16 than in the average (π=6.3×10^−3^) of 18 maize genes in previous reports (Ching et al., 2002). This finding implied weak or no natural selection pressure on ZmbHLH16 and provided evidence that ZmbHLH16 is highly evolutionarily conserved in maize. The target gene sequence polymorphism also reflects evolutionary pressure during maize improvement (Wang et al., 2005). Previous studies found one polymorphic site per 60.8 bp in maize (Ching et al., 2002). In the present experiment, a lower frequency was obtained for ZmbHLH16 in 78 maize inbred lines (one SNP or InDel every 69.8 bp). The global LD decay of ZmbHLH16 investigated in our study (*r^2^*<0.1 within 1,300 bp) was also less than the average intragenic level (*r^2^*<0.1 within 1,500 bp) (Remington et al., 2001). The above nucleotide polymorphism testing results confirmed the conserved evolution of ZmbHLH16. The conserved molecular evolution of ZmbHLH16 further hinted at its crucial function in maize male reproduction.

Most bHLH proteins consist of a classical helix-loop-helix (HLH) domain to form homo- or heterodimers with other HLH-proteins to regulate downstream target genes (Murre et al., 1989). bHLH-bHLH or bHLH-MYB complexes have been reported to be involved in plant fertility (Niu et al., 2013; Qi et al., 2015; Chen et al., 2016). Our experiments showed that ZmbHLH16 lacks the ability of transcriptional activation. Thus, we speculate that ZmbHLH16 might regulate target gene expression by interacting with other proteins. One of its interacting factors, ZmbHLH51, was identified and confirmed using genome-wide coexpression and yeast two-hybrid analyses. In rice, the BIF domain is necessary for DYT1-bHLH protein dimerization (Cui et al., 2016). The present study showed that the BIF domain is also present in ZmbHLH16(D) ^241–365 a.a.^ and participates in the interaction between ZmbHLH16 and ZmbHLH51. Previous studies related to bHLH proteins mainly focused on their conserved domains. In the investigation of the interaction between ZmbHLH16 and ZmbHLH51, we noticed that not only the conserved bHLH and BIF domains but also the ZmbHLH16(B) ^81–160 a.a.^. region could form heterodimers with ZmbHLH51. Moreover, the ZmbHLH16(G) ^161–365 a.a.^ fragment may have a negative effect on activation, leading to a reduced transcription activation capacity of the full-length ZmbHLH16 protein. Taken together, our findings provide new evidence that in bHLH proteins, other regions are of importance in molecular function in addition to the typical bHLH and BIF domains.

Coincidently, it was recently reported that ZmbHLH16 was a candidate gene for the ms23 mutant (Nan et al., 2017). The tapetal layer of the ms23 mutant undergoes abnormal periclinal division instead of tapetal differentiation (Chaubal et al., 2000). These results strongly support the hypothesis that ZmbHLH16 is involved in tapetal specification and maturation. In Nan’s paper (Nan et al., 2017), the researchers mainly focused on the abortion mechanism in the *ms23* mutant, combining RNA-seq with proteomics data. These authors also confirmed the interaction between ZmbHLH16 and ZmbHLH51, similar to our result. In contrast, we paid more attention to the ZmbHLH16 nucleotide polymorphisms, molecular evolution, expression features, subcellular location and regulatory mechanisms. Through coexpression analysis, a group of genes potentially involved in maize male reproduction were also revealed in this study. Our results might help uncover the mechanism of ZmbHLH16 regulating the pollen abortion in the ms23 mutant.

## Acknowledgments

We thank the Chinese Maize Industry Technology System for providing maize inbred lines in this experiment with the aid of Prof. Guangtang Pan and Prof. Lujiang Li. We sincerely thank Prof. Shibin Gao for providing the genomic DNA of several maize inbred lines. We also thank Prof. Yufeng Hu for providing the original vectors. We are very grateful to Dr. Yibing Yuan and Jingtao Qu for assisting in data analysis.

## Footnotes

1 http://www.maizesequence.org

2 http://www.linux1.sαftberry.cαm/berry,phtml?tαp¡c=fgenesh&grαup=prαgrams&subgrαup=gf¡nd

3 http://www.ncb¡.nIm.n¡h.gov/Structure/cdd/wrpsb.cg¡

4 http://www.gramene.org/

5 http://qteller.com/ma¡ze2/

6 http://202.120.45.92/addb/

7 ftp://ftp.ensemblgenomes.org/pub/plants/release-29/fasta/zea_mays

8 http://b¡o¡nfo.cau.edu.cn/agr¡GO/analysis.php

9 http://www.omicshare.com/tools

